# A Deep Redox Proteome Profiling Workflow and Its Application to Skeletal Muscle of a Duchene Muscular Dystrophy Model

**DOI:** 10.1101/2022.08.15.504013

**Authors:** Nicholas J. Day, Tong Zhang, Matthew J. Gaffrey, Rui Zhao, Thomas L. Fillmore, Ronald J. Moore, George G. Rodney, Wei-Jun Qian

## Abstract

Perturbation to the redox state accompanies many diseases and its effects are viewed through oxidation of biomolecules, including proteins, lipids, and nucleic acids. The thiol groups of protein cysteine residues undergo an array of redox post-translational modifications (PTMs) that are important for regulation of protein and pathway function. To better understand what proteins are redox regulated following a perturbation, it is important to be able to comprehensively profile protein thiol oxidation at the proteome level. Herein, we report a deep redox proteome profiling workflow and demonstrate its application in measuring the changes in thiol oxidation along with global protein expression in skeletal muscle from *mdx* mice, a model of Duchenne Muscular Dystrophy (DMD). In depth coverage of the thiol proteome was achieved with >18,000 Cys sites from 5608 proteins in muscle being quantified. Compared to the control group, *mdx* mice exhibit markedly increased thiol oxidation, where ~2% shift in the median oxidation occupancy was observed. Pathway analysis for the redox data revealed that coagulation system and immune-related pathways were among the most susceptible to increased thiol oxidation in *mdx* mice, whereas protein abundance changes were more enriched in pathways associated with bioenergetics. This study illustrates the importance of deep redox profiling in gaining a greater insight into oxidative stress regulation and pathways/processes being perturbed in an oxidizing environment.

**Graphical Abstract:** 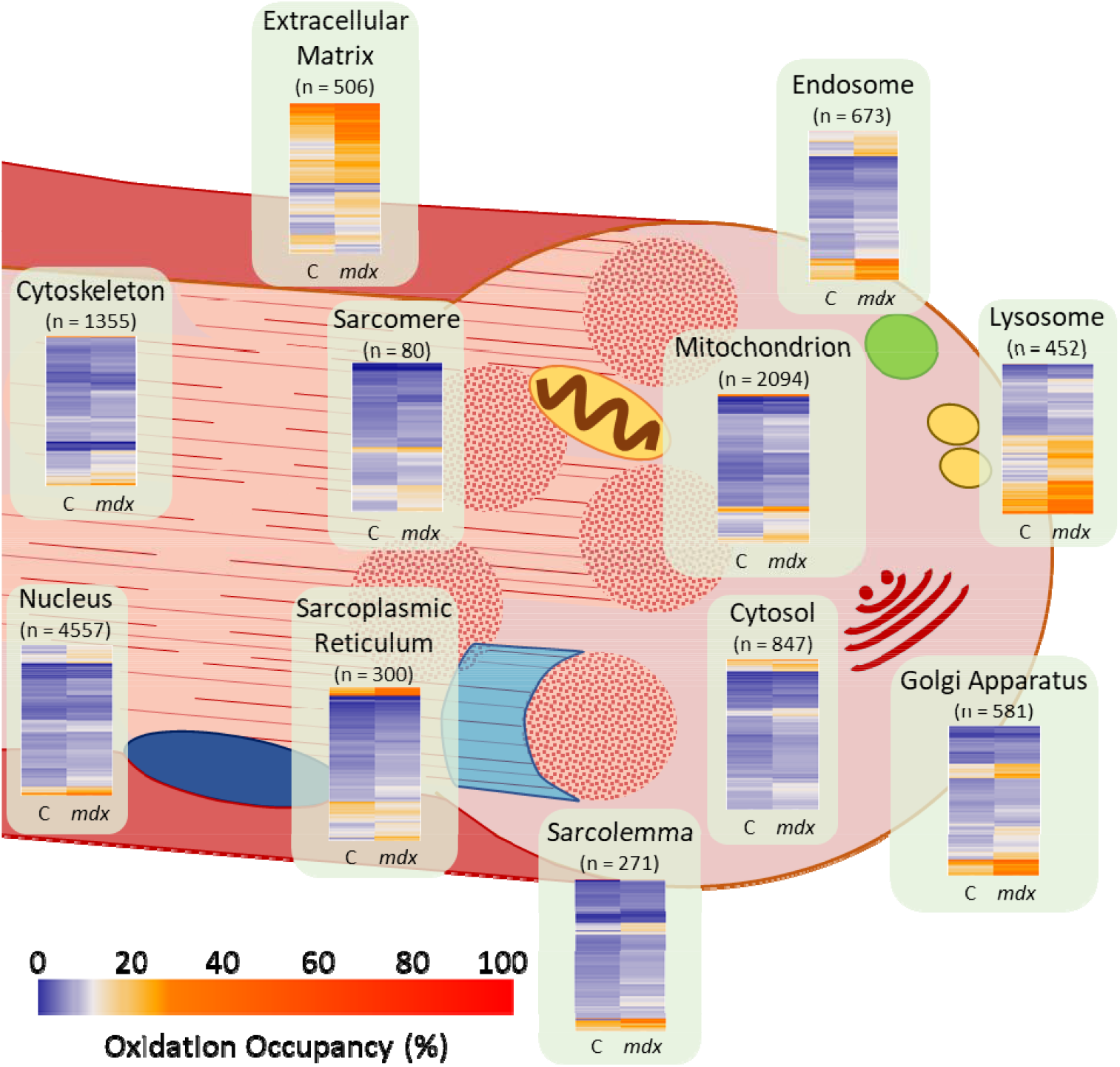

**Highlights:**

- Deep redox profiling workflow results in stoichiometric quantification of thiol oxidation for > 18,000 Cys sites in muscle
- Thiol redox changes were much more pronounced than protein abundance changes for the overlapping set of proteins
- Redox changes are most significant in coagulation and immune response pathways while abundance changes on bioenergetics pathways

## 1. Introduction

Redox regulation is fundamental for both normal cellular activities and disease development. Redox signaling under physiological conditions, defined as oxidative eustress [1], is an important mechanism for many underlying processes such as metabolism [2], immune response [3], and proliferation [4]. Redox homeostasis is maintained by balancing the level of cellular oxidants such as reactive oxygen/nitrogen/sulfur species (ROS/RNS/RSS) and antioxidants, predominantly in the form of enzymes or small molecules [5, 6]. An imbalanced redox system that favors the oxidants leads to oxidative distress (or stress) and dysregulated of redox signaling [7]. Oxidative stress accompanies many diseases and response mechanisms, such as cancer [8], neurodegeneration [9], metabolic disorders [10], pathogen clearance [11], exposure to nanoparticles [12], and becomes more prominent with aging [13].

A major mechanism of redox regulation is through post-translational modifications (PTMs) of the cysteine thiol group of proteins, which often act as “redox switches”. Protein thiol-based modifications can induce significant modulatory changes to protein structure, catalysis, protein-protein interactions, and/or binding capacity [14–17]. The free cysteine thiol can be oxidized to a range of different reversible forms, including disulfides (S-S), S-sulfenylation (SOH), S-nitrosylation (SNO), S-glutathionylation (SSG), S-persulfidation (SSH), and S-polysulfidation (SS_n_H) [18]. Further oxidation of SOH leads to the formation of S-sulfinylation (SO_2_H) and S-sulfonylation (SO_3_H), which are generally considered as irreversible modifications and can serve as indicators of prolonged oxidative stress [19]. Despite reports of S-sulfinylation being reduced back to free thiols by sulfiredoxin [20, 21], the occurrence of this mechanism is limited. Thus, most studies have focused on reversible thiol-based modifications.

Reversible thiol redox PTMs enable Cys sites to function like “switches” to regulate pathway activity or induce the activity of an alternate pathway [17, 22–25], much like phosphorylation [26]. Thiol redox switches can also activate “moonlighting” properties of multi-functional proteins, causing them to execute an alternate function when modified [27, 28]. Furthermore, our group has recently demonstrated that Cys sites are occupied by multiple types of PTMs [29, 30], emphasizing the importance of PTM diversity in regulating outcomes of pathways or protein function. The occurrence and occupancy of these modifications partially depend on the reactivity of the thiol group, which is further determined by the microenvironment of specific cysteine residues [18, 31]. Other key factors include genetic mutations and external stimuli. As a result, modulatory redox PTMs under homeostatic oxidative eustress conditions can lead to microscopic effects on the overall global redox state, while strong oxidative stress could result in macroscopic effects on the global redox state [32].

Given the complexity and the broad scope of redox regulation across the proteome[33], it is highly important to have analytical technologies that comprehensively profile Cys sites subject to redox modifications in the proteome (redox proteome) and quantify the site-specific occupancy at large scale. However, quantitative characterization of the thiol redox proteome with in-depth coverage has been a challenge to date despite significant advances in redox proteomics [34–36]. Current methods still do not provide sufficient coverage of the thiol proteome. This is largely due to the low-abundance nature of proteins bearing redox PTMs and the stochastic sampling approach used in current mass spectrometry-based proteomics methods. Finally, deep profiling of the redox proteome can be even more difficult in challenging tissues such as skeletal muscle because of the presence of dominant proteins, leading to the quantification of only ~4,500 Cys sites at most, as reported in a recent study [35].

Herein we report a deep redox profiling workflow for greatly enhancing the coverage of thiol redox proteome and multiplexed stoichiometric quantification of thiol oxidation in tissues (or cells). The present workflow builds upon our recently reported RAC-TMT (Resin-Assisted Capture coupled with Tandem Mass Tags) approach [30, 37] by incorporating a microscale fractionation scheme to enhance the proteome coverage. In addition, our workflow can also profile protein abundance quantification of the global proteome, thus allowing the differentiation between redox and protein abundance changes. To demonstrate the utility of this approach, skeletal muscle from a mouse model of Duchenne Muscular Dystrophy (DMD) (*mdx* mutant mice with truncated dystrophin) were analyzed and compared to control mice.

Oxidative stress is known to be involved in the pathogenesis of myopathies [38] and muscular diseases such as Duchenne muscular dystrophy (DMD), a condition caused by X chromosome-linked mutations in the dystrophin gene, have a birth prevalence of 20 per 100,000 males [39]. Molecularly, dystrophin forms a dystrophin-glycoprotein complex that links the intracellular cytoskeletal network of the sarcolemma to the extracellular matrix and is essential for providing strength, contraction, and stability in skeletal muscle [40, 41]. By contrast, dysfunctional dystrophin in DMD patients causes sarcolemma damage, which further leads to a range of secondary effects including inflammation, impaired regeneration, and oxidative stress [42, 43]. Sources for elevated ROS production in DMD patients have also been identified, including NADPH oxidases (NOX), nitric oxide synthase (NOS), and xanthine oxidase (XO) [42, 44]. In addition, perturbed Ca^2+^ homeostasis has been proposed as a signaling paradigm in mediating redox-related pathophysiology of DMD [45, 46]. The *mdx* mouse is the most widely used animal model for DMD research. Despite being deficient for dystrophin, the *mdx* mice have been reported with minimal clinical symptoms and their lifespan is only reduced by ~25% compared to control mice {McGreevy, 2015 #106}. In line with this notion, our deep redox profiling revealed strong upregulation of thiol oxidation in the *mdx* mice compared to control and broad effects of DMD on an array of biological processes.

## 2. Materials and Methods

### 2.1 Animal care and tissue collection

Control (C57BL/10SnJ, Stock No:000666) and *mdx* (C57BL/10ScSn-Dmd^mdx^/J, Stock No:001801) were purchased from The Jackson Laboratory, housed in a specific pathogen free facility on a 12hr light/dark cycle. Male mice 17-24 weeks of age were deeply anesthetized by isoflurane (2%) inhalation and euthanized by rapid cervical dislocation in accordance with National Institutes of Health guidelines and approved by the Institutional Animal Care and Use Committee of Baylor College of Medicine. Tibialis anterior muscles were dissected and rapidly frozen in liquid N2 and stored at −80°C until preparation for redox proteomics.

### 2.2 Synthesis of Thiol Affinity Resin

The original Thiopropyl Sepharose 6B used to capture protein thiols is discontinued by the manufacturer and we have developed a protocol to synthesize the resin in-house using similar chemistry as in the original resin. The in-house resin is prepared by coupling *N*-hydroxysuccinimide (NHS)-Activated Sepharose 4 (GE Healthcare) with 2-(Pyridyldithio) ethylamine hydrochloride (MedChem Express). Note: all centrifugation steps are performed at 100 x g for 2 min. Briefly, 5 mL of suspended resin slurry was transferred to a 10 mL spin column and centrifugated to remove the isopropanol. The resin was thoroughly washed 5X with 5 mL wash buffer (1mM HCl) and centrifugated. The resin was then washed once with 5 mL of coupling buffer (0.2M NaHCO3, 0.5M NaCl, pH 8.3) and centrifugated. Next, 175 mg of 2-(Pyridyldithio) ethylamine hydrochloride was dissolved with 3.5mL of coupling buffer. 3mL of dissolved 2-(Pyridyldithio) ethylamine hydrochloride was then added directly to the 1.5g of drained NHS-activated resin. The reaction column was supported by placing it in a 50 mL tube and was incubated for 2 hours at room temperature with shaking at 1000 RPM in a thermomixer. Following the reaction, the column was centrifugated and the resin was washed 5X with 5 mL coupling buffer. To block unreacted NHS groups on the resin, 3mL of quenching buffer (1M Tris pH 7.5) was then directly added to the drained resin and incubated at room temperature for 1 hour with shaking at 1000 RPM in a thermomixer. The resin was then washed 5X with 5 mL ultrapure water, 3X with 3 mL isopropanol. Finally, enough isopropanol was added to resuspend the resin in a final volume of 7.5 mL and final concentration of 200 mg/mL. Resin stock stored at 4°C is usable up to 9 months from the date of synthesis.

### 2.3 Proteomic sample preparation

Sample preparation was comprised of parallel workflows for quantifying protein abundance, total thiol oxidation, and total thiol content. For total oxidation, frozen tissues were first minced into small pieces on an aluminum tray with dry ice. For each sample, around 20 mg tissue was incubated in lysis buffer (250 mM MES, pH 6.0, 1% SDS, 1% Triton-X-100) with 100 mM N-ethyl-maleimide (NEM) for 30 min and protected from light. Two additional samples were generated by pooling tissue from either the control or *mdx* genotype for quantification of “total thiol”, including both reduced and oxidized protein thiols (**Fig. 1A**). Total thiol samples were prepared in a similar manner as total oxidation samples except without NEM in the lysis buffer. Tissue homogenization was performed with a Tissue-Tearor homogenizer (BioSpec Products). The lysate was cleared by centrifugation and proteins were precipitated with cold acetone overnight at −20°C. The pellet was washed with acetone and then dissolved in resuspension buffer (250 mM HEPES, pH 7.0, 8 M urea, and 0.1% SDS). The protein concentrations were determined based on the bicinchoninic acid assay (BCA, Thermo Fisher Scientific). At this point, the total thiol samples were split into three technical replicates, for a total of 3 total thiol channels per genotype **(Fig. 1B**). Buffer exchange of a 500µg aliquot of each sample was performed with 8 M urea in 25 mM HEPES, pH 7.0 using a 0.5 mL 10K Amicon ultra filter (MilliporeSigma). The proteins were reduced with 20 mM dithiothreitol (DTT) at 37□ for 30 min. Excessive reducing reagents were removed by buffer exchange.

**Figure 1.**
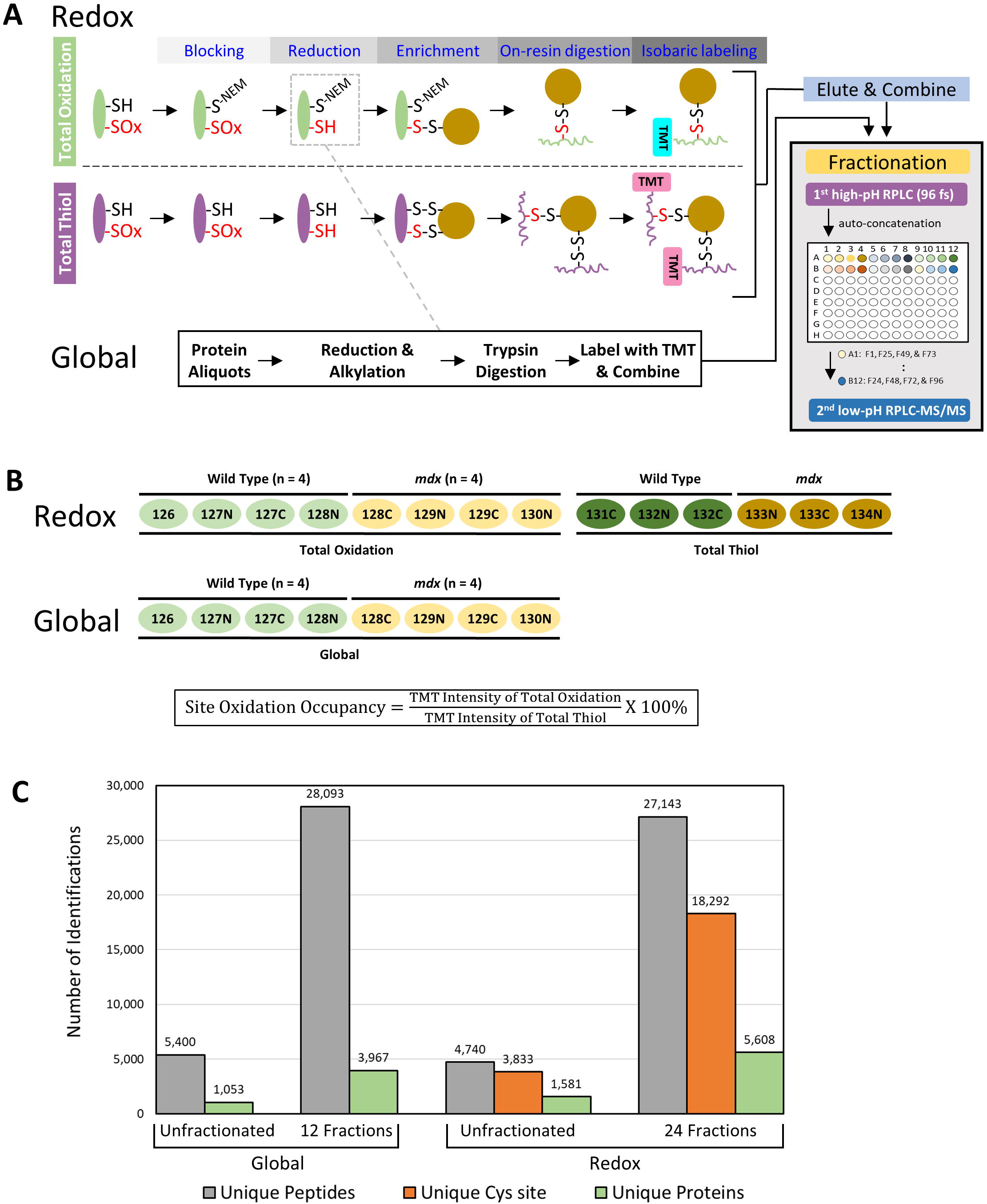
The deep redox and global profiling workflow. (A) Schematic of the workflow including sample preparation and LC-MS/MS. All forms of reversible oxidation are represented by a red “-SOx” group. Parallel processing of “oxidation” and “total thiol” samples consists of initial blocking of *in situ* protein free thiols with NEM (omitted during the total thiol sample preparation), reduction of reversibly oxidized cysteine thiols by DTT, enrichment via resin-assisted capture, and on-resin digestion and isobaric TMT labeling. Samples for global proteomics are collected following the reduction step of the total oxidation workflow (gray dashed box) and are processed in parallel (boxed workflow below total thiol schematic). The combined eluted samples are then separated into 96 fractions by off-line high-pH RPLC and concatenated into 12 (global) or 24 (redox; illustrated in A) fractions with each fraction being analyzed by LC-MS/MS. (B) TMT labeling schemes for the redox and global proteomics samples in this study, along with the formula for calculating the site occupancy of thiol oxidation. (C) Bar plot showing the coverage of global and redox proteomes with or without fractionation. Colored bars represent the number of unique peptides (gray), unique Cys sites (orange), or unique proteins (green).

100µg aliquots of protein were collected from each total oxidation sample (total thiol channels excluded) for global protein abundance profiling. Briefly, the proteins were dissolved in 40 µL 25mM HEPES pH 7.7, reduced with 5 mM DTT at 37°C for 30 min., and alkylated with 10mM iodoacetamide at room temperature for 1 hour. The proteins were digested with Trypsin (1 µg/µL) at 37°C overnight, labeled with TMT following the manufacturer’s instructions, quenched with 5% hydroxylamine, combined, cleaned up by C18 SPE (solid phase extraction) (Phenomenex).

The RAC-TMT workflow was performed similarly as previously described [47] using the in-house synthesized thiol-affinity resin. Briefly, 30 mg of resin is used to capture 150µg of reduced protein with nascent free thiols for each channel. Following capture, proteins are digested on-resin and the remaining peptides are labeled with TMT and eluted by DTT. TMT16 regents were used for multiplexing of enriched samples **(Fig. 1B)**. 130C and 131N were not used for total oxidation quantification due to the potential interference from the 131C and 132N tags that were used to quantify the total thiol. Global protein abundance samples that correspond with total oxidation samples followed the same labeling scheme.

Following elution and combination of enriched total oxidation peptides as previously described [36, 47], the eluents were alkylated with iodoacetamide at a concentration that is 4-fold greater than the concentration of DTT present in the sample from the elution. All enriched samples within the same TMT set were combined, desalted using SPE.

### 2.4 High-pH reverse phase capillary LC fractionation

Peptides from both the redox and global procedure were fractionated using a microscale LC fractionation setup as previously described [29, 48]. Briefly, 30µg of peptides were fractionated using a reversed-phase LC column (65 cm × 200 µM internal diameter (ID) packed with 3μm Phenomenex Jupiter C18 particles) on a nanoAcquity LC system (Waters). The binary solvent buffers (~pH 7) are comprised of mobile phase A (10mM ammonium formate in water) and mobile phase B (acetonitrile). Eluted peptides were separated into 96 fractions in 1-minute intervals, which were concatenated into 24 or 12 fractions (**Fig. S1**) using an autosampler (HTC PAL; CTC analytics) running Chronos software. The flow rate was kept at 2.2 µL/min. Fractions were dispensed into wells of a 96 well plate containing 20 µL of 0.03% N-Dodecyl-β-maltoside to facilitate collection and prevent peptide loss. Samples were stored frozen until final LC-MS/MS analysis.

### 2.5 LC-MS/MS

LC-MS/MS was performed as previously described [49]. Briefly, for each fraction, the peptides were analyzed on a Q Exactive Plus mass spectrometer (Thermo Scientific) with a nano-electrospray ion source. The peptides were separated on a self-packed reverse-phase column (60 cm x 50 µm ID with Jupiter 3 μm C18 material (Phenomenex)) with an integrated PicoTip emitter (New Objective) using a 2 hr LC gradient. A data-dependent acquisition method was used with the following settings. Full MS scans (m/z 400-1,800) was collected at a resolution of 70,000 with maximum ion injection time of 20 ms and automatic gain control of 3E6. Subsequent fragment ion spectra were collected for up to 12 most abundant precursor ions. Precursor ions were first selected (isolation width 2 m/z) and fragmented by higher collisional energy-induced dissociation (HCD) at 32% normalized collisional energy (NCE). MS/MS scans were acquired at a resolution of 35,000 with maximum ion injection time of 200 ms and automatic gain control of 1E5. Fixed first mass was set to 110 m/z to include TMT reporter ions and dynamic exclusion was set to 30 sec.

### 2.6 Data analysis

LC-MS/MS data were searched using MS-GF+ [50] against *Mus musculus* protein sequences from Uniprot (Release 2019-9). Key parameters include a parent ion mass tolerance of 20 ppm, partial tryptic rule with up to 2 missed cleavages. Oxidation on methionine (+15.9949), NEM blocking of Cysteine (+125.047679), and Carbamidomethylation of Cysteine (+57.0215) were selected as dynamic modifications. TMT labels on peptide N-termini and lysine (+304.207146) were selected as fixed modifications. Peptide spectra matches (PSMs) were filtered using the following criteria: 1) mass accuracy within 10 ppm; 2) PepQ value < 0.01 to a final false-discovery rate < 1%. The TMT reporter ion intensities were extracted by MASIC [51]. For global or enriched samples, data from all fractions were combined.

For global protein abundance data, peptide TMT reporter ion intensities were aggregated to the unique protein level by summing up raw reporter ion intensities (non-log2 transformed) of corresponding peptides or PSMs. For data normalization, reporter ion intensities of all channels were log2 transformed and normalized by median-centering. For redox data, TMT reporter ion intensities of all channels were log2 transformed and total oxidation channels within each genetic group were median-centered due to the significant differences in redox levels observed between two genotypes. Total thiol channels were normalized by median-centering across groups. To calculate the thiol oxidation stoichiometry for each Cys site, the data was aggregated from the unique peptide level to the unique Cys site level using an in-house R script. The Cys site stoichiometry was then calculated as the difference (in log2 scale) between the intensity of total oxidation for a given sample and the average intensity of the corresponding total thiol samples.

Statistical analysis was performed using a Student’s t test with Benjamini & Hochberg (BH) method for false discovery control. To investigate the coverage for transcription factors and cofactors, we obtained a list of known transcription factors and cofactors from AnimalTFDB 3.0 [52]. Pathway analysis was performed by the use of core analysis with default settings by Ingenuity Pathway Analysis (IPA) (QIAGEN Inc.) [53]. For global abundance data, proteins were filtered by adjusted p value < 0.05. For redox data, only one Cys site was assigned to represent its corresponding protein. For proteins where multiple Cys sites were identified, the Cys site with highest log2FC value was selected for further analysis. The filtering criteria for the enrichment data first involved sorting by p adjusted value < 0.05, where the absolute log2FC value of Cys sites was used to determine the cutoff value for the top 2 quartiles (i.e. median) in the dataset. After the data was filtered by absolute log2FC > top2 quartile cutoff, the Cys sites must have a % occupancy that is greater than 10% in both the control and *mdx* backgrounds. Display of protein structures and Cys sites was performed using ICM-Browser Pro.

### 2.7 Data availability

The raw datasets presented in this study will be deposited in online repositories.

## 3. Results

### 3.1 Assessment of the deep redox profiling workflow

To enhance the coverage of the thiol redox proteome, we modified the RAC-TMT workflow (**Fig. 1A**), which was previously developed in our lab [36, 47], by incorporating a microscale fractionation scheme into the workflow. Briefly, for profiling thiol total oxidation (i.e., all reversible forms of thiol PTMs), protein free thiols are initially blocked with NEM, reversible thiol oxidation is then reduced with DTT, and proteins with nascent free thiols are enriched using a thiol-affinity resin. The captured proteins are then digested on-resin with trypsin and the peptides are labeled on-resin with TMT (**Fig. 1A**; see “Total Oxidation”). For preparation of “total thiol” samples in parallel, the initial NEM blocking step was omitted (**Fig. 1A**; see “Total Thiol”). Following TMT-labeling, the eluted peptides from all channels were combined and fractionated by microscale LC into 24 concatenated fractions (**Fig. S1**) with each fraction being analyzed by LC-MS/MS, thus enhancing the overall thiol proteome coverage. Within this workflow, we also take protein aliquots from each total oxidation sample after the DTT reduction step (**Fig. 1A**; see “Global”), to profile the global proteome in terms of protein abundance.

In this case, muscle tissue from 4 control and 4 *mdx* mutant mice were processed using a total oxidation workflow and total thiol samples were generated by pooling tissue of all 4 samples per genotype using 16-plex TMT reagents. Each genotype’s total thiol sample was split into 3 technical replicates during processing (**Fig. 1B**). Two channels (130C and 131N) were omitted to avoid “crosstalk”[54] between total thiol and total oxidation TMT channels. Including multiple total thiol channels in our workflow enables robust stoichiometric measurement of thiol oxidation occupancy (**Fig. 1B**). These total thiol channels provide an additional benefit, as they ensure that enough enriched thiol-containing peptides are present for better detection by mass spectrometry, a concept often referred to as “boosting”, as described in a previous phosphoproteomics study [55]. The increased amounts of total thiol-containing peptides contributed from total thiol channels also make the workflow more amenable to fractionation, since the reversibly oxidized Cys-containing peptides by themselves are often insufficient for effective fractionation. In this case, the inclusion of total thiol channels in the sample multiplex design is critical for providing enough peptide to make offline high pH reverse-phase LC (RPLC) fractionation feasible (e.g., > 10µg) [56], as total thiol channels are expected to contain more enriched peptides than total oxidation channels.

**Fig. 1C** shows that fractionation of both redox and global samples significantly improves overall coverage compared to unfractionated samples. Fractionation of the redox sample led to identification of more than 27,000 unique peptides, which amount to ~18,300 Cys sites quantified, corresponding to 5,608 proteins. This observed coverage of the thiol proteome is nearly 4 times the number of quantified Cys sites reported in the recent OxiMouse study[35] for muscle. The deep coverage of thiol redox proteome is also nicely illustrated by the drastic improved coverage of transcription factors and cofactors (**Fig. S2**), an important class of low-abundance proteins that are typically challenging to detect. Consequently, the enhanced coverage in our study allows us to investigate both thiol oxidation and protein abundance alterations between control and *mdx* muscle tissues in greater detail.

### 3.2 Deep redox profiling reveals a widespread increase in thiol oxidation in *mdx* muscle

The inclusion of multiple total thiol channels per genotype in our multiplexing scheme permitted the calculation of the stoichiometry of oxidation (site occupancy) for each biological sample (**Fig. 1B**), which was then used for calculation of stoichiometric fold change and statistical analysis (**Supplemental Data 2**). Stoichiometric quantification was highly reproducible, as the CVs for this data were approximately 5% (**Fig. S3A**). Plotting the total oxidation occupancy of all Cys sites identified in our study (18,292) revealed a median increase of ~2% in overall thiol oxidation in *mdx* mice compared to control mice (**Fig. 2A**), suggesting that the redox state in *mdx* mice is perturbed towards a more oxidative environment. Subcellular localization analysis of corresponding proteins revealed that Cys sites exhibiting greater than 20% oxidation in *mdx* mice belong to proteins in multiple subcellular compartments, such as the extracellular matrix, lysosome, endosome, golgi, and sarcoplasmic reticulum (**Fig. 2B**).

**Figure 2.**
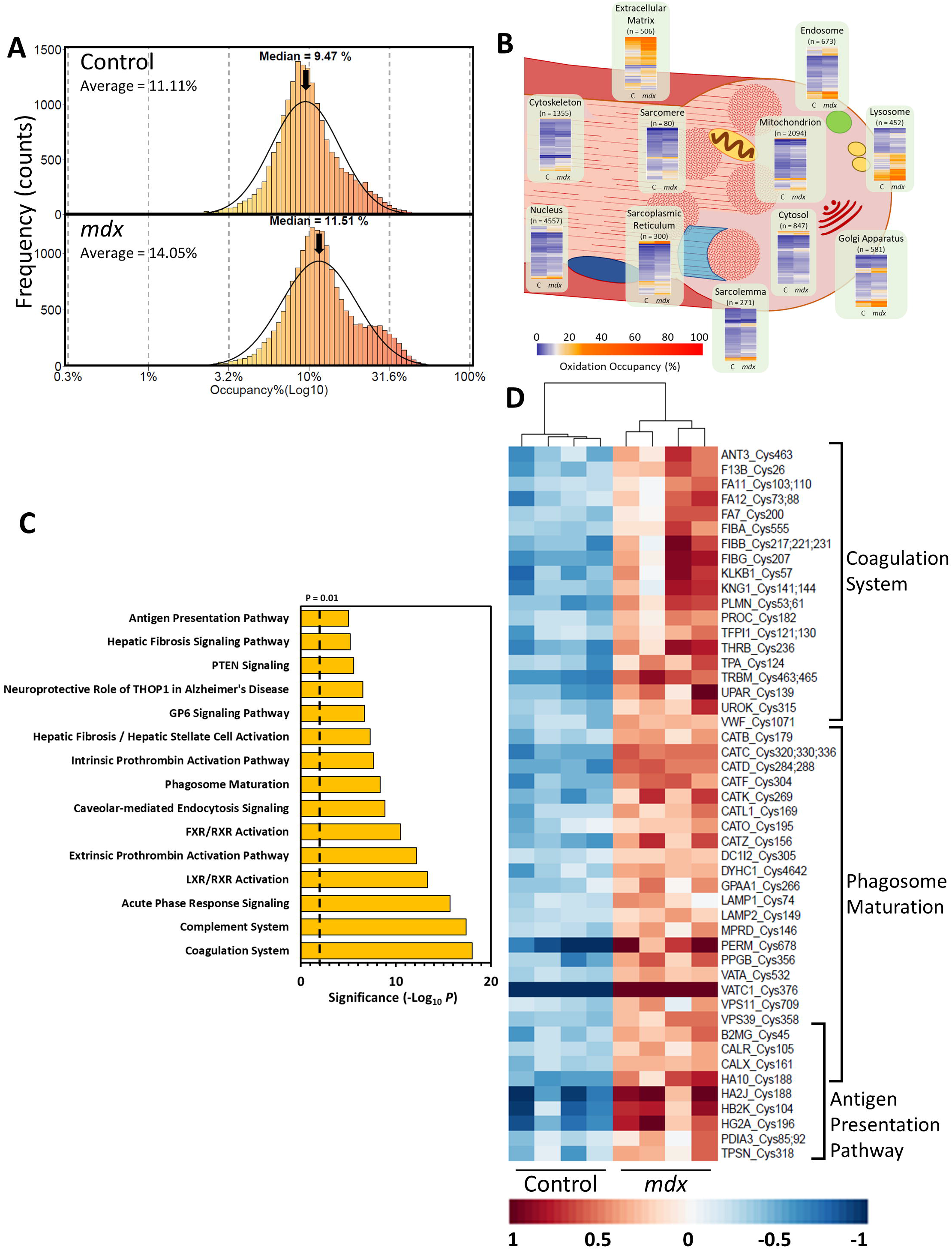
The *mdx* mutation exerts a strong perturbation on the thiol redox proteome. (A) Histogram representing the distribution of total oxidation occupancies at the Cys site level, which is plotted on a log10 scale for control and *mdx* mice. The curve represents the normal distribution of Cys sites in the histogram, where the density is scaled to the frequency (y axis). (B) Muscle cell cartoon showing heatmaps of Cys site oxidation occupancies in control (C) or *mdx* mice. Cys sites were assigned to specific subcellular compartments based on their corresponding protein’s uniprot annotation. The number of Cys sites (n) for each location is reported below each compartment category. (C) Bar plot representing the top 15 most enriched categories recovered from IPA for proteins with Cys sites that exhibited significant changes in oxidation. The cutoff for category significance (p < 0.01) on the −log10 transformed x axis is denoted by the dashed line. (D) Representative heatmap of proteins belonging to “Coagulation System”, “Phagosome Maturation”, or “Antigen Presentation Pathway” categories. Overlapping brackets for “Phagosome Maturation” and “Antigen Presentation” denote proteins shared by these categories. For each row (i.e., a protein’s most oxidized Cys site), the mean centered stoichiometry per replicate is plotted, where the mean represents the average stoichiometry across all replicates for both genotypes.

To learn more about what canonical pathways are potentially impacted by the redox alterations, we performed IPA analysis for those proteins with significant redox changes. Since we are interested in proteins and Cys sites that displayed substantial alterations in redox levels in *mdx* mice, we specified criteria for identifying significantly altered proteins for IPA analysis (see Methods). Briefly, for the 18,292 Cys sites, each protein was assigned with the most altered Cys site based on log2FC values to represent the protein (**Fig. S3C**; n = 5,608). Subsequent filtering required these protein Cys sites to have: an adjusted p value (padj) < 0.05, a log2FC within the top 2 quartiles, and finally the average site occupancy must be greater than 10% (**Fig. S3C**; n = 888; **Supplemental Data 3**). A volcano plot of these proteins confirms our earlier observation that the data is enriched in proteins and corresponding Cys sites that exhibit increased oxidation (884 of 888 total) in the *mdx* mutant mice (**Fig. S3C**). Only 4 of 888 significant proteins had Cys sites with decreased oxidation (RPC10, PLM, DOC10, and PARVG), where these sites were the only ones detected for these proteins.

### 3.3 A Diverse set of pathways are associated with increased thiol oxidation

A closer investigation of the 888 significant proteins and their Cys sites with predominantly increased oxidation via IPA revealed that these proteins belong to an array of pathways (**Supplemental Data 4**). The most significantly enriched pathway was “Coagulation System” (**Fig. 2C**), which is accompanied by related pathways such as “Extrinsic [or Intrinsic] Prothrombin Activation Pathway” as shown in the pathway overlap network (**Fig. S3D**). 19 proteins from our dataset comprise the “Coagulation System” category and many of these are factors that are necessary to form blood clots, such as FIBB/FIBG (Fibrinogen beta/ gamma chain), VWF (Von Willebrand Factor), PLMN (Plasminogen), THRB (Prothrombin), and TRBM (Thrombomodulin) (**Fig. 2D**). Our redox data also hints at potential perturbation of the procoagulant “GP6 Signaling Pathway” category (**Fig. 2C**). We observed increased oxidation of collagen VI (CO6A1/2) in our data, a protein important for blood clot formation and the microfilament network of the extracellular matrix [57, 58]. These results suggest that oxidative stress may be a factor in the antagonism between pro- and anti-coagulation mechanisms and could contribute to DMD pathology.

Immune response related pathways also appear to be significantly enriched with the oxidation data, as “Complement System” and “Acute Phase Response Signaling” were ranked the next two most significant categories after “Coagulation System” (**Fig. 2C**). Proteins belonging to “Acute Phase Response Signaling” overlap with 7 other pathways (**Fig. S3D**), illustrating a potential redox-dependent regulation mechanism that encompasses multiple pathways. Further examination of the Cys sites and proteins corresponding to “Phagosome Maturation” and “Antigen Presentation” (**Fig. 2D**) revealed how thiol oxidation could perturb other aspects of immunological processes. Notably, subunit C1 of the V-type H^+^ ATPase proton pump (VATC1) and numerous thiol proteases (cathepsins; PPGB and CAT[B,C,D,F,K,L1,O,Z]) exhibited increased thiol oxidation in *mdx* mice (“Phagosome Maturation”; **Fig. 2D**). In the context of phagosomes, proton pumping helps to lower the pH of the phagosome to activate low pH proteases (such as cathepsins) for degradation of internalized material like pathogens or debris from apoptotic cells [59, 60]. This suggests that thiol oxidation of components critical for the efficacy of lysosomal/phagosome-degradation may also contribute to autophagic dysfunction observed in *mdx* [61, 62]. As for antigen presentation, which depends on phagosome uptake of debris or pathogens [63], we observed several proteins belonging to the major histocompatibility class I or II complexes (MHC) (HA10, HA2J, HB2K, and HG2A) that exhibited increased oxidation in *mdx* mice (**Fig. 2D**). Our results may provide insight into other ways adaptive immune system signaling is modulated in DMD.

### 3.4 Drastic shifts in thiol oxidation occupancy on a variety of proteins

The high coverage afforded by our deep redox profiling approach allows for recovery of more potentially biologically relevant Cys sites. As shown in **Fig. 2A**, some Cys sites in *mdx* mice show oxidation occupancies greater than 20%, creating the appearance of a secondary bump in the *mdx* histogram, potentially highlighting functional Cys sites in response to oxidative stress. This prompted us to further investigate these sites at the structural level. We applied a stringent threshold comprised of: log2FC > ±1 and an absolute difference in oxidation occupancy (Δocc) between *mdx* and control mice that is greater than ±20%, which identified 51 sites with marked changes in oxidation (**Supplemental Data 5**). Using Δocc as a metric allowed for more stringent filtering, as log2FC values are not always representative of dramatic shifts in thiol oxidation compared to stoichiometric/occupancy measurements (i.e., log2FC = 1 could be a 2 to 4% or 20 to 40% shift). The heatmap in **Fig. 3A** illustrates the contrast between genotypes for these sites with large shifts in occupancy.

**Figure 3.**
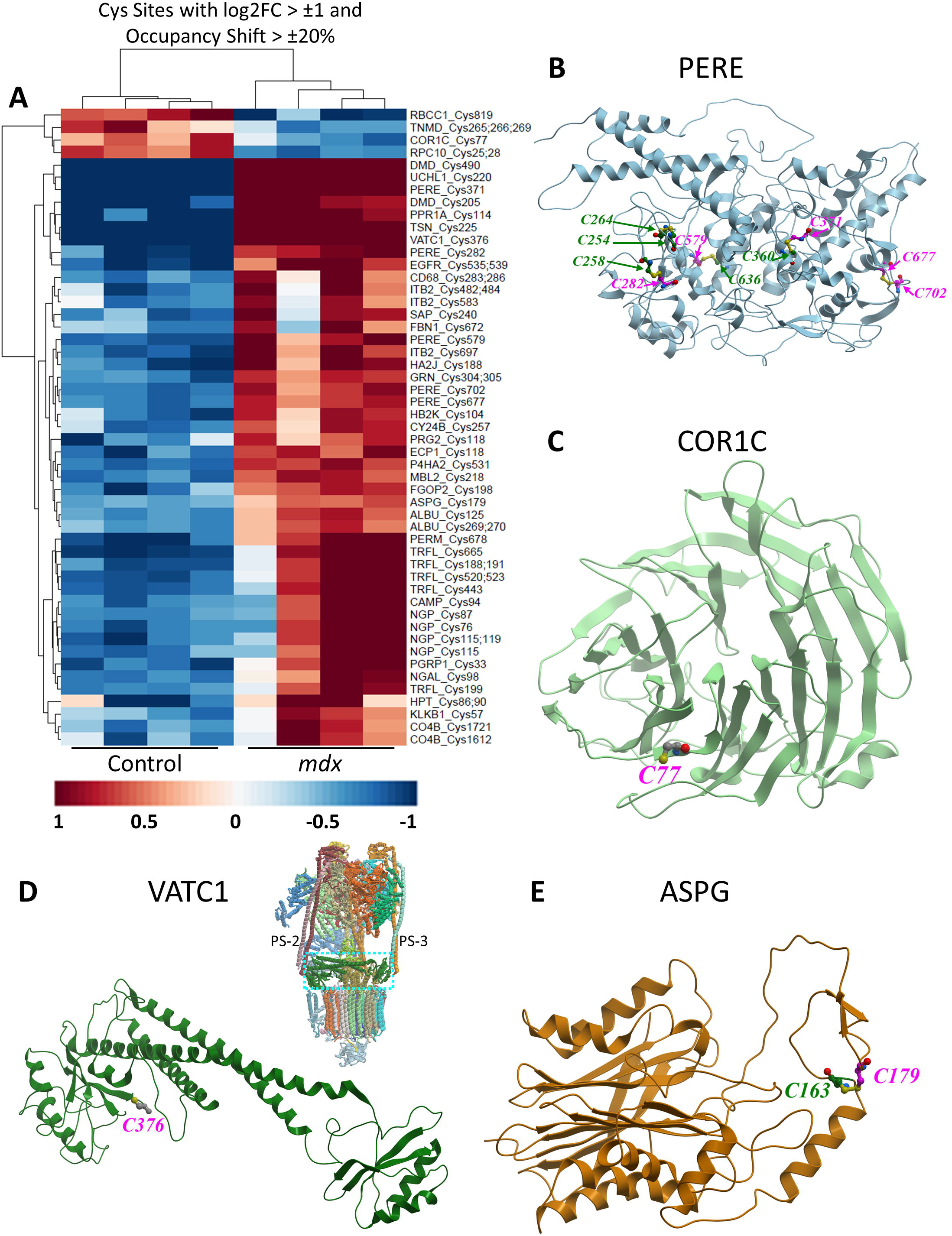
Structures of proteins with highly oxidized Cys sites. (A) Representative heatmap of Cys sites with oxidation stoichiometries that meet the criteria of log2FC > ±1 and a difference in absolute oxidation occupancy (Δocc) that is greater than ±20% (also found in **Supplemental Data 5**). Each row represents a Cys site with its mean-centered stoichiometries plotted, where the mean represents the average stoichiometry across all replicates for both genotypes. (B-E) Structures of proteins with Cys sites exhibiting dramatic changes in oxidation as identified in (A) and **Supplemental Data 5**, are highlighted with magenta colored text. Neighboring Cys sites (insignificantly oxidized or not in the data) are also highlighted for reference in green colored text. (B) Mouse eosinophil peroxidase (PERE). Cys 282 is pictured in a disulfide with neighboring Cys 258, where either Cys could also potentially participate in disulfide formation with Cys254 or 264. Other potential disulfides include Cys360-371, 579-636, and 677-702. (C) Human coronin-1C (COR1C) (PDB: 7STY). (D) Human V-type ATPase complex (PDB: 6wm4) [68], where the dotted cyan box denotes the location of the C1 subunit (VATC1) shown in the adjacent structure. PS = peripheral stalk. (E) N(4)-(beta-N-acetylglucosaminyl)-L-asparaginase (ASPG). Note: The AlphaFold protein structure prediction program [64] was used to display the structures for PERE (AlphaFold identifier: AF-P49290-F1) and ASPG (AlphaFold identifier: AF-Q64191-F1).

Among the sites with large shifts in oxidation was Cys220 of the ubiquitin C-terminal hydrolase L1 (UCHL1), which we identified in a previous study as sensitive to ER stress-induced oxidation in pancreatic beta cells [29]. Eosinophil peroxidase (PERE) showed significant increases in oxidation in *mdx* mice at 5 sites (Cys 282, 371, 579, 677, and 702), with Cys 371 and 282 showing more than a 40% increase (**Supplemental Data 5**). A closer look at the AlphaFold-predicted structure [64] of PERE reveals that these Cys sites may form disulfides with neighboring Cys residues (**Fig. 4B**). These disulfides likely promote structural stability and/or folding, however more work is needed to understand their impact on the function of this enzyme. Coronin-1C (COR1C) was an interesting observation, as Cys 77 exhibited an approximately 43% reduction in oxidation in *mdx* mice compared to the control. Visualization of COR1C shows that Cys 77 is part of a flexible loop belonging to blade one of the seven blades that make the β-propellor architecture of the protein (**Fig. 4C**) [65]. COR1C binds to and crosslinks filamentous actin [66] as well as promotes trafficking of Rac1 for fibroblast migration [67]. Future work is needed to understand the significance of Cys 77 oxidation, or lack thereof, on these processes.

**Figure 4.**
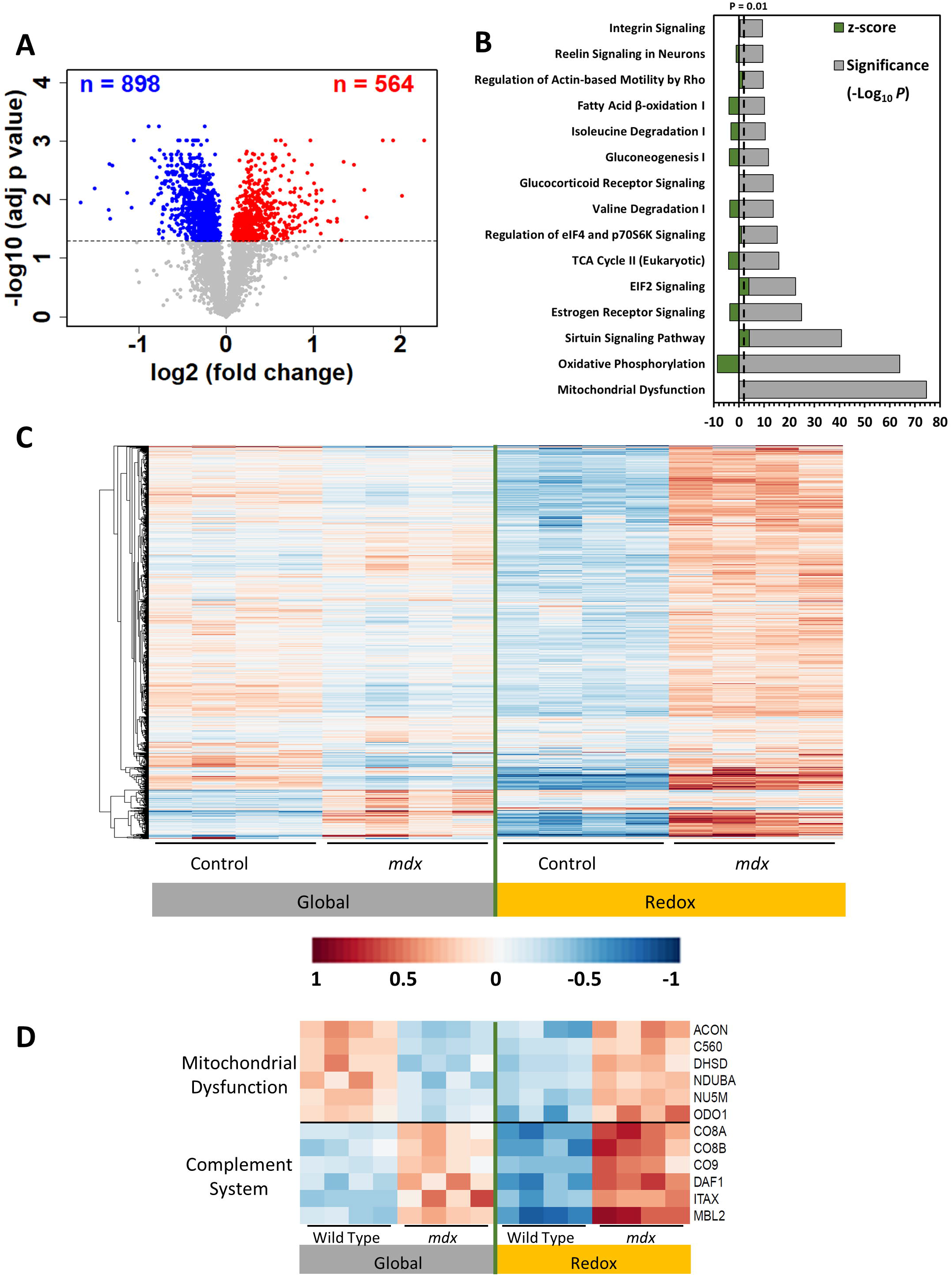
Protein abundance changes in *mdx* mice and its contrast to redox changes. (A) Volcano plot representing adjusted p values (on −log10 scale) and log2 fold changes in protein abundance between control and *mdx* mutant mice. Gray data points represent proteins that do not meet the significance cutoff of padj < 0.05, while red and blue data points denote proteins that are significantly up- or down-regulated, respectively. (B) Bar plot representing the top 15 most enriched categories recovered from IPA for proteins with significant changes in protein abundance. The cutoff for category significance (p < 0.01) on the −log10 transformed x axis is denoted by the dashed line. Z-score bars (green) are plotted over the p value bars (gray), which represent the activation state of the pathways (positive value indicates upregulation, while negative values indicate downregulation). Note that no Z scores were generated by IPA for the “Mitochondrial Dysfunction” and “Glucocorticoid Signaling” categories. (C) Heatmap plotting the 3,107 proteins common to both the global and redox enrichment datasets, where each row represents a single protein. For the global data, the normalized TMT intensities of each replicate were mean-centered, while the stoichiometries of each replicate corresponding to the Cys site with the greatest log2FC value were mean-centered and plotted to represent the same protein for the redox data. Note that the global and redox data were mean centered independently and then merged to generate the heatmap. (D) Representative heatmap of proteins corresponding to “Mitochondrial Dysfunction” or “Complement System” pathways identified in both redox and global datasets (**Supplemental Data 4**). Data for the heatmap was prepared and utilized in the same manner as with panel C. The heatmap scale in panel C also applies to the heatmaps in panel D.

VATC1 Cys 376 is another interesting example of a significant difference in oxidation (**Fig. 4D**), with *mdx* mice showing a ~36% increase in oxidation compared to the control (**Supplemental Data 5**). The tertiary structure of VATC1 is comprised of two α-helices that link together “head” and “foot” domains, which interact with coiled coils named peripheral stalks (PS) 2 and 3, to form a collar that stabilizes the quaternary structure of the V-type ATPase (**Fig. 4D**) [68]. Cys 376 is in the “foot” domain that interacts with PS2, where oxidation at this site could perturb the assembly or stability of the ATPase complex, which uses ATP to actively transport 3H^+^ and increase luminal H^+^ levels of multiple types of organelles, including phagosomes [69]. Also related to phagosome-lysosome function is N(4)-(beta-N-acetylglucosaminyl)-L-asparaginase (ASPG), also known as lysosomal aspartylglucosaminidase (AGA), which is a hydrolase that breaks down glycoproteins in lysosomes. Deficient activity of ASPG results in the lysosomal storage disease aspartylglycosaminuria [70], which is caused by a Cys163Ser mutation that prevents formation of a disulfide bridge, leading to its inability to be properly folded and activated [71, 72]. Equally, a Cys179Ser mutation has a similar effect on ASPG folding, leading to its rapid degradation [72]. We observed a nearly 32% increase in oxidation at ASPG Cys 179 in *mdx* mice, which is known to form disulfides with Cys 163 (**Fig. 4E**) based on a previous report [73]. This Cys 163-179 disulfide is critical for local loop structure stability in the α-subunit, as it promotes contact with the β-subunit to form a heterodimer [74], which later forms a heterotetramer [75]. In summary, these studies emphasize the importance of measuring Cys site oxidation to identify sites that could be determinants of protein stability, assembly, and function.

### 3.5 Redox changes were more pronounced compared to protein abundance changes

Global proteome profiling is complementary to redox profiling, which allows us to compare redox level changes with protein abundance changes. In this workflow, we profiled protein abundance in parallel from the same set of starting samples, where more than 28,000 unique peptides, which correspond to 3,967 unique proteins were identified (**Fig. 1C, Supplemental Data 1**). The coverage is comparable to a recent human muscle tissue proteomic study that also implemented offline fractionation [76]. Compared to the control, the *mdx* genotype leads to substantial changes in protein abundances (**Fig. 4A**). High reproducibility of these quantification was observed (**Fig. S4A**). P-value distribution histograms indicated that the abundances of many proteins were significantly different between the genotypes (**Fig. S4B**). With adjusted p (padj) < 0.05, 1,462 proteins were observed with significant changes (**Fig. 4A**).

To obtain a better understanding of the pathways involved with the significant changes in abundance for more than 1400 proteins, similar IPA analysis was performed for canonical pathways (**Supplemental Data 4**). **Fig. 4B** shows the top ranked pathways enriched in the dataset, including “Mitochondrial Dysfunction”, “Oxidative Phosphorylation”, “TCA Cycle (Eukaryotic)”, “Gluconeogenesis”, “Valine (or Isoleucine) Degradation”, and “Fatty Acid β-oxidation”. Our dataset provides high coverage of proteins corresponding to all five complexes of the mitochondrial electron transport chain (as represented “Mitochondrial Dysfunction” schematic in **Fig. S4C**) and almost all proteins related to these processes have significantly reduced abundance in *mdx* mice. The z-scores assigned by IPA for these pathways represent their predicted activation state, which are negative for these categories, reflecting a strong downregulation of bioenergetics in *mdx* mice based on protein abundance (**Fig. 4B**). This aligns with other proteomics reports that have identified reduced abundance of mitochondrial proteins in *mdx* muscle tissue [77–79].

Interesting, for proteins identified in both the redox and global datasets, the redox changes (stoichiometry of oxidation) were observed to be much more pronounced than the protein abundance (translational) changes (**Fig. 4C)**, supporting the significant role of redox regulation in oxidative stress conditions. **Fig. 4D** further illustrates the distinctive regulation at the translational and posttranslational levels. For mitochondrial dysfunction, all proteins were down-regulated in their expression in *mdx*, but more oxidized in *mdx*. The decreased protein expression and increased oxidation are likely to work synergistically, leading to mitochondrial dysfunctional. In complement system, both the protein expression and oxidation are increased in *mdx*, but oxidation is more substantial compared to abundance changes. Together, these findings highlight the dynamic nature of both translational and post-translational regulation in *mdx* muscle and suggest that they can function independently under normal or perturbed conditions.

## 4. Discussion

The present study demonstrates a deep redox profiling workflow for achieving multiplexed quantification with high coverage of the thiol proteome along with global proteome profiling. The utility of this approach was well demonstrated in the application of muscle from the DMD mouse model. The strong induction of oxidative stress in *mdx* mice compared to control mice is reflected by a strong distribution of proteins with significantly altered Cys site oxidation. The stoichiometric measurements of the thiol redox proteome show that the *mdx* genotype is strongly perturbed by the oxidative stress that is characteristic of DMD and impacts proteins belonging to a variety of different pathways.

The redox proteomics approach implemented in this study enabled us to deeply profile the thiol redox proteome of muscle tissue to a much greater level than any prior redox proteomic study, including the recent Oximouse report [35]. A key element in our workflow was the implementation of offline fractionation, which significantly improved coverage. The obtained coverage is comparable to what can be obtained in a typical phosphoproteomic study [80] and is unprecedented for redox proteomics, given the challenge of profiling muscle due to the presence of dominant proteins [81]. Furthermore, the multiple replicates of genotype-specific total thiol channels, which are representative of all free and reversibly oxidized thiols present in the sample, made quantification of thiol oxidation at the stoichiometric level possible. Stoichiometry-based analysis offers an advantage towards understanding changes in thiol oxidation[32]. There are also several caveats to consider for the current approach. Perhaps the major caveat for the thiol oxidation profiling is that we do not have information on the specific types of PTMs, especially since the thiol PTM landscape has been reported as very complex [29]. It will be necessary to integrate thiol oxidation profiling with other approaches if specific PTM information is desired. Also, the achievable coverage of the thiol proteome is likely dependent on the number of fractions being collected and analyzed.

We note that the integration of redox proteome and global proteome profiling for the DMD model also offers an interesting case study of translational and posttranslational regulation. In our data, we have identified a wide range of proteins and Cys sites that are perturbed in *mdx* mice. Interpreting the significant changes in abundance and/or Cys site oxidation is important to improving our understanding of the pathophysiology of DMD.

Our global data points to a strong disruption of bioenergetic-related processes as a prominent theme in *mdx* mice. Downregulated expression of proteins in pathways such as “TCA Cycle (Eukaryotic)”, “Gluconeogenesis”, “Valine (or Isoleucine) Degradation”, and “Fatty Acid β-oxidation” (**Fig. 4B**), suggests metabolic dysfunction as an accompanying component of DMD pathology, which has been proposed by others [82, 83]. Significant representation of downregulated proteins associated with “Mitochondrial Dysfunction” or “Oxidative Phosphorylation” pathways (**Fig. 4B**) supports metabolic dysfunction, thereby emphasizing the importance of mitochondrial homeostasis in DMD [82, 84]. The observed reduction of ATP production in *mdx* muscle tissue [85, 86] could in part be due to the reduced expression of proteins belonging to each of the electron transport chain complexes (**Fig. S4C**). Previous reports indicate that mitochondrial dysfunction precedes both skeletal and cardiac muscle impairment in DMD [87–89], making it an area of continued focus for future DMD-related studies.

In contrast, the redox data point to perturbation of a different set of biological processes, with coagulation and immune response pathways being the most notable. Intriguingly, coagulation and acute phase response processes are proposed to originate from a common mechanism [90], as they may share stimuli [91] and proteins, such as thrombin, which can cleave pro-IL1α to active IL1α for induction of inflammation [92]. The protein overlap map supports this proposal, as “Acute Phase Response Signaling” overlaps with “Coagulation System” and thrombin activation pathways (**Fig. S3D**). Abnormalities or disorders in blood coagulation are occasional side-effects of DMD pathology [93, 94], but the molecular mechanisms behind this remain unclear. Oxidation of proteins in the “Complement System” is another avenue by which immunological processes are potentially perturbed. As complement C3 is critical for initiation of immune responses [95, 96], its oxidation may influence downstream processes, such as “Phagosome Maturation” and “Antigen Presentation Pathway” (**Fig. 2C**). Since autophagy is vital for homeostasis of muscle tissue[97], pathologies with an increased demand for regeneration following repeated injuries, like DMD, succumb to muscle degeneration when autophagy is impaired [61, 62, 98]. Consequently, oxidation poses a challenge to this essential process. Previous reports have shown that autophagy impairment results from increased oxidative stress in DMD [62, 98] or even the loss of Collagen VI[99], which is important for coagulation and is part of the“GP6 Signaling Pathway”. In line with this, we observed increased oxidation of Collagen VI (CO6A1/2) in our data, which may be relevant to perturbation of both the coagulation system as well as phagosome maturation and proper autophagy.

Enrichment of the “PTEN Signaling” category (**Fig. 2C**) may also be relevant to the perturbed autophagy theme. A recent study showed in *mdx* mouse myoblasts that increased phosphatidylinositol-3,4,5-trisphosphate 3-phosphatase (PTEN) signaling led to upregulated autophagosome formation but yielded a reduced overall flux [100], suggesting defective autophagy. It is worth noting that our group has since identified increased Nox2-derived ROS and decreased acetylation of α-tubulin as unrelated disruptors of autophagosome-lysosome fusion in *mdx* mice (under review [101]), suggesting that multiple perturbed pathways may converge to cause impaired autophagy. Together, these findings suggest that increased oxidative stress in *mdx* mice perturbs pathways involved in blood clotting and immune response, but also potentially has a strong effect on autophagic mechanisms that are critical for muscle regeneration.

## 5. Conclusion

The deep redox profiling workflow demonstrated in this study provides unprecedented coverage and a comprehensive view of the thiol redox and global proteomes that are substantially perturbed by the *mdx* mutation in muscle tissue. In general, the redox changes appear to be more pronounced than protein abundance changes. Significant changes in protein abundance are most enriched in pathways associated with bioenergetics, while marked changes in thiol oxidation are observed in proteins associated with immune response signaling. Proteins in these majorly perturbed pathways are shared with other pathways, suggesting that the *mdx* mutation induces a complex perturbation that is part of the pathogenesis of the disease.

## Abbreviations

DMD: Duchenne Muscular Dystrophy
IPA: Ingenuity pathway analysis
NEM: N-ethyl-maleimide
NOS: Nitric oxide synthase
PTM: Post-translational modification
RAC: Resin-assisted capture
RAC-TMT: Resin-Assisted Capture coupled with Tandem Mass Tags
RNS: Reactive nitrogen species
ROS: Reactive oxygen species
RPLC: Reverse phase liquid chromatography
RSS: Reactive sulfur species
SDS: Sodium dodecyl sulfate
SOH: S-sulfenylation
SO_2_H: S-sulfinylation
SO_3_H: S-sulfonylation
SSG: S-glutathionylation
SSH: S-persulfidation
SS_n_H: S-polysulfidation
S-S: Disulfide

## Acknowledgements

The proteomics work was performed in the Environmental Molecular Sciences Laboratory, Pacific Northwest National Laboratory, a national scientific user facility sponsored by the Department of Energy under Contract DE-AC05-76RL0 1830.

## Figure Legends

**Supplemental Figure S1**. Off-line peptide fractionation by high-pH reversed phase liquid chromatography. Schematics of auto-concatenation for consolidation of 96 fractions into 24 or 12 fractions, where four or eight non-consecutive fractions are pooled into one concatenated fraction, respectively.

**Supplemental Figure S2**. Deep redox profiling enables greater coverage of transcription factor oxidation. Comparison of transcription factor (A) or transcription cofactor (B) coverage in samples with or without fractionation. In A and B, colored bars represent the number of unique peptides (gray), unique cysteine residues (orange), and unique proteins (green), respectively. (C) Comparison of oxidation occupancies for 296 transcription factor Cys sites in control and *mdx* mice. (D) Comparison of oxidation occupancies for 1088 transcription cofactor Cys sites in control and *mdx* mice. Diagonal red dashed lines in C-D indicate y = x. Colors indicate the number (Count) of neighboring data points to represent density.

**Supplemental Figure S3**. Quantification metrics of the redox proteomics dataset. (A) Quantification reproducibility, determined by coefficient of variation (CV), within biological replicates. The dashed line represents a CV threshold of 10%. (B) Histograms of raw (left) or BH-adjusted p values (right) representing comparisons between control and *mdx* mice. The blue column denotes how many p values are less than 0.01, while the dashed red line indicates the adjusted p value cutoff of 0.05. (C) Volcano plots representing adjusted p values (on −log 10 scale) and log2 fold changes in Cys site oxidation between control and *mdx* mutant mice of Cys sites meeting stringent criteria for IPA (see section 3.2 in main text). *Left*: Plotting each protein with its most oxidized Cys site. *Right:* All the proteins that meet the subsequent criteria (padj < 0.05, log2FC > top 2 quartile, % occupancy > 10%). (D) Overlap network of the IPA redox data categories shown in **Fig. 2C**. Each line between different IPA pathway nodes indicates overlap of proteins between those pathways, with the number (positioned approximately midway on the line) representing how many proteins overlap. A minimum of 5 overlapping proteins was needed to plot a pathway.

**Supplemental Figure S4**. Quantification metrics of the global proteomics dataset. (A) Quantification reproducibility, determined by coefficient of variation (CV), within biological replicates. The dashed line represents a CV threshold of 10%. (B) Histograms of raw (left) or BH-adjusted p values (right) representing comparisons between control and *mdx* mice. The blue column denotes how many p values are less than 0.01, while the dashed red line indicates the adjusted p value cutoff of 0.05. (C) Schematic (derived from IPA) of proteins corresponding to the category “Mitochondrial Dysfunction”. Proteins outlined in purple were detected in the significant proteins dataset and protein names are in SwissProt format. Note the high representation of mitochondrial proteins in the electron transport chain (complexes I-IV), where some of these proteins were shown in a representative heatmap in **Fig. 4D**. (D) Overlap network of the IPA global data categories shown in **Fig. 4B**. Each line between different IPA pathway nodes indicates overlap of proteins between those pathways, with the number (positioned approximately midway on the line) representing how many proteins overlap. A minimum of 5 overlapping proteins was needed to plot a pathway.

