## Supplementary figures and images for "A Deep Redox Proteome Profiling Workflow and Its Application to Skeletal Muscle of a Duchene Muscular Dystrophy Model"

### Supplemental Figures

Supplemental Figure S1

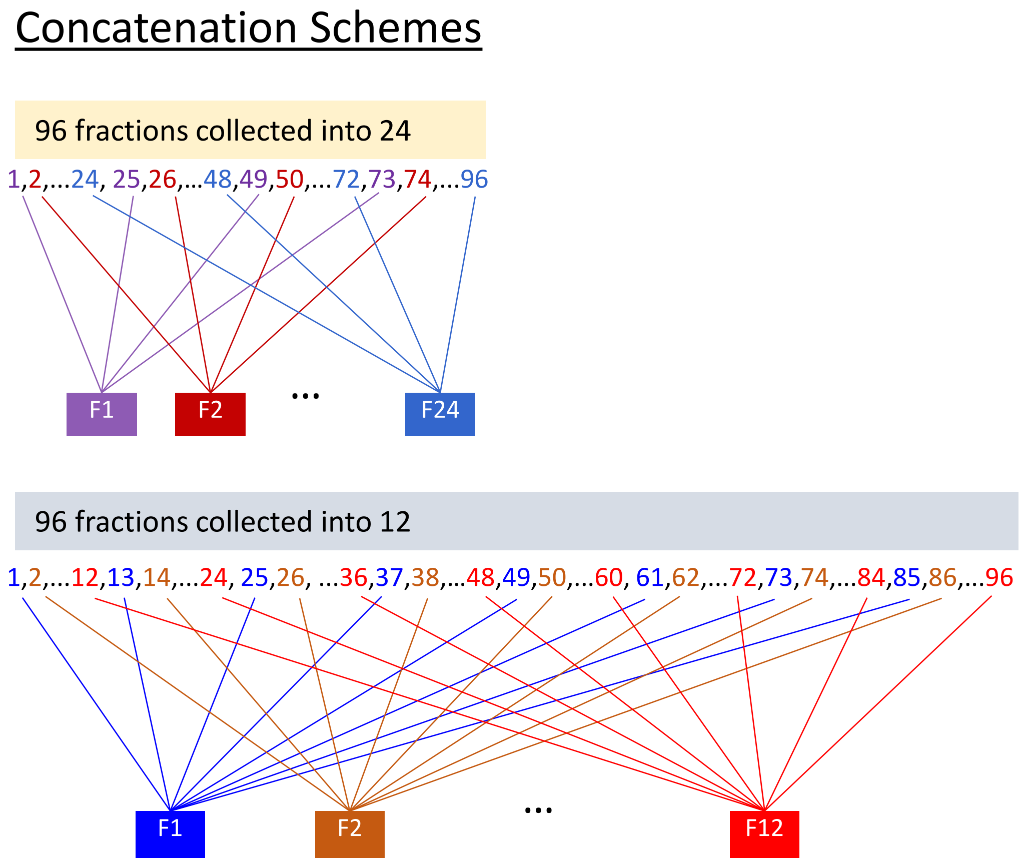

Supplemental Figure S2

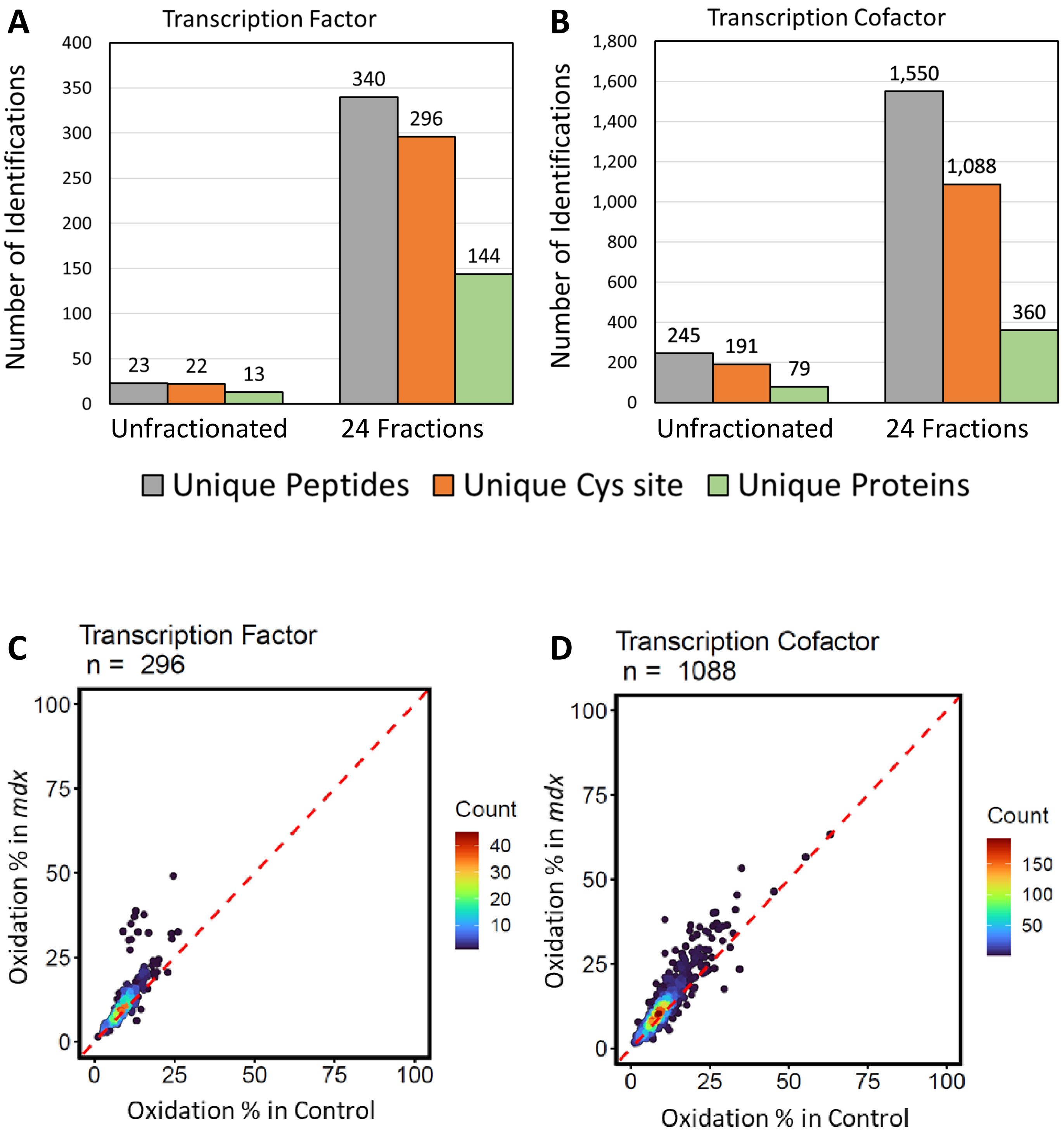

Supplemental Figure S3

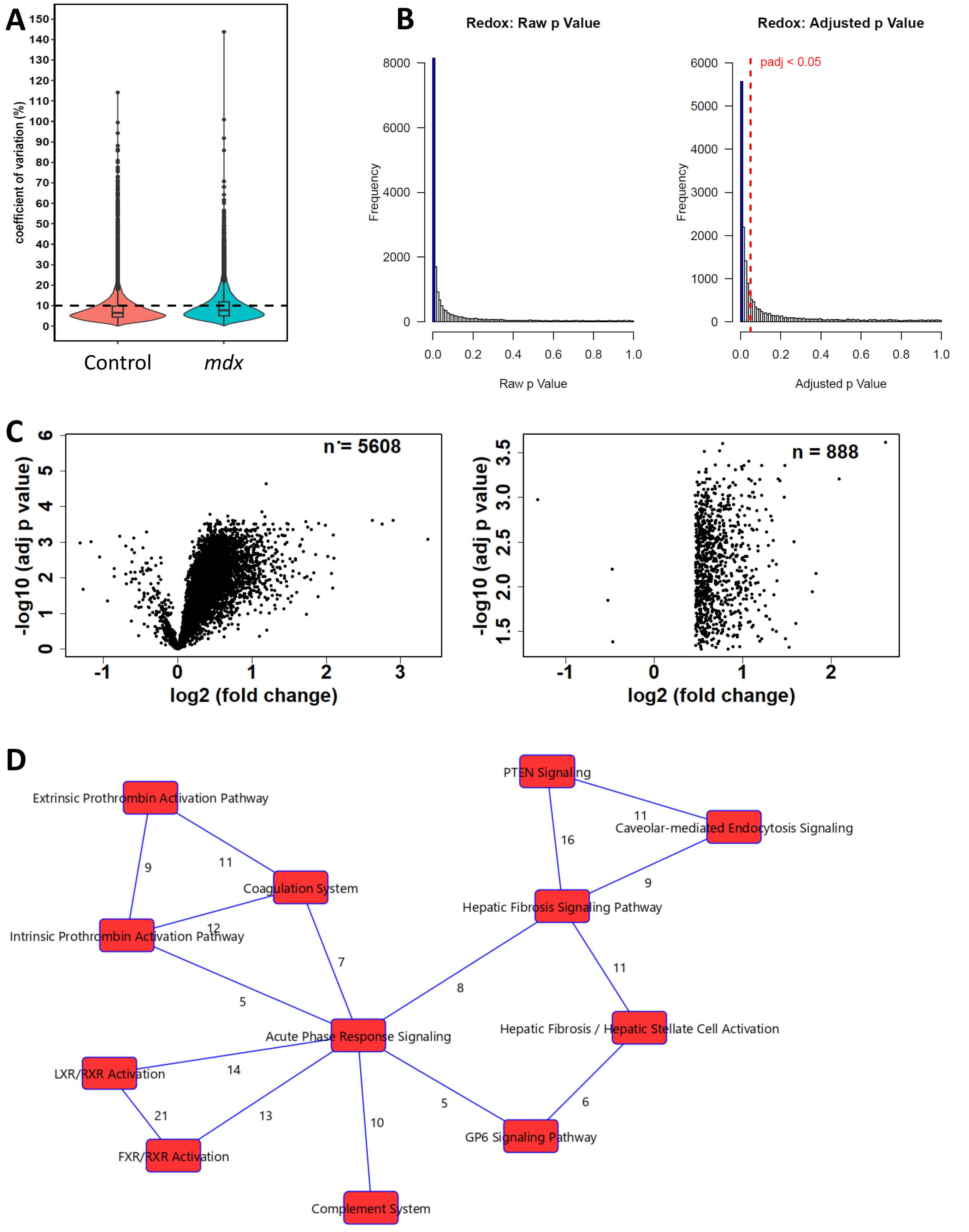

#### Supplemental Figure S4

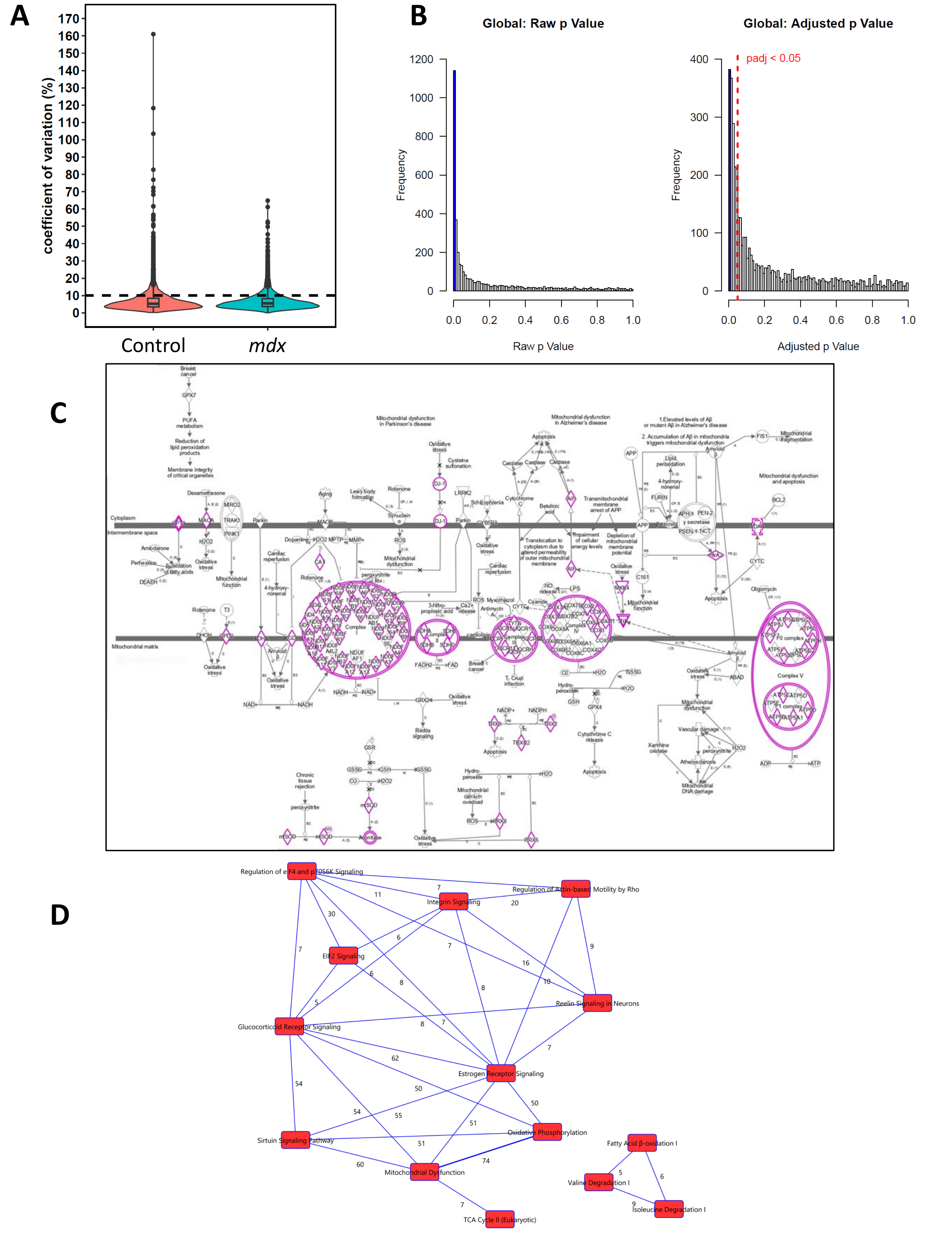
